# Alteration of the chicken upper respiratory microbiota, following H9N2 avian influenza virus infection

**DOI:** 10.1101/2023.08.08.549695

**Authors:** Tara Davis, Dagmara Bialy, Joy Leng, Roberto La Ragione, Holly Shelton, Klaudia Chrzastek

## Abstract

Several studies have highlighted the importance of the gut microbiota in developing immunity against viral infections in chickens. We have previously shown that H9N2 avian influenza A virus (AIV) infection retards the diversity of the natural colon associated microbiota, which may further influence chicken health following recovery from infection. The effects of influenza infection on the upper respiratory tract (URT) microbiota are largely unknown. Here we showed that H9N2 AIV infection lowers alpha diversity indices in the acute phase of infection in URT, largely due to the family Lactobacillaceae being highly enriched during this time in the respiratory microbiota. Interestingly, microbiota diversity did not return to levels similar to control chickens in the recovery phase after viral shedding has ceased. Beta diversity followed similar trend following challenge. *Lactobacillus* associate statistically with the disturbed microbiota of infected chickens at acute and recovery phase of infection. Additionally, we study age-related changes in the respiratory microbiota during maturation in chickens. From 7 to 28 days of age, species richness and evenness were observed to advance over time as the microbial composition evolved.

Maintaining microbiota homeostasis might be considered as a potential therapeutic target to prevent or aid recovery from H9N2 AIV infection.

## Introduction

Avian influenza A viruses (AIVs) belong to the Orthomyxoviridae virus family and have a single- stranded, negative sense RNA genome (Spackman, 2020). AIVs can be categorised into high and low pathogenicity viruses (HPAIVs and LPAIVs, respectively), depending on their disease severity (Alexander, 2007). HPAIVs have up to 100% mortality rates and infect chickens systematically, whereas LPAIVs are associated with milder symptoms and lower mortality rates and are limited to respiratory and intestinal tracts (Bottcher-Friebertshauser et al., 2013). This difference in pathogenicity is determined by the haemagglutinin (HA) protein, based on which proteases recognise the proteolytic cleavage site sequence (Spackman, 2020). The H9N2 subtype was first identified in China in the early 1990s and is now considered to be endemic in poultry throughout Asia, Europe, North Africa and the Middle East (Li et al., 2017, Sun and Liu, 2015, Peacock et al., 2019). Despite the H9N2 subtype generally having a mortality rate of less than 20%, outbreaks are associated with decreased egg production and clinical symptoms, as well as an increased likelihood of secondary infections with potentially higher mortality rates (Gu et al., 2017, Lee et al., 2016). It is also considered zoonotic, as it can be transferred to humans and can also be transferred to other mammals, such as pigs (Mostafa et al., 2018, Gu et al., 2017). Therefore, being able to treat and control the spread of H9N2 AIV in poultry populations is important for preventing economic losses and protecting public health.

There is growing evidence that the commensal bacteria present at the body sites of chickens play an important role in shaping the host’s defences against viral infection (Abaidullah et al., 2019). Variations in the gut microbiota composition have been shown to be caused by viral infection, as well as environmental factors like housing, diet and hygiene (Kers et al., 2018, Clavijo and Florez, 2018). The relationship between the gut microbiota and AIV in the chicken model has been observed in numerous studies (Yitbarek et al., 2018a, Yitbarek et al., 2018b, Yitbarek et al., 2018c, Li et al., 2018). These studies have shown that H9N2 AIV infection is associated with the dysregulation of the gut microbiota, as well as changes in innate immune gene expression levels. Yitbarek et al. showed that the depletion of the gut microbiota by a cocktail of antibiotics leads to a significant downregulation of the interleukin-22 (IL-22) (Yitbarek et al., 2018b). IL-22 is an important cytokine in the gastrointestinal tract, as it plays a role in maintaining homeostatic levels and host defences against microbes, particularly in barrier repair (Keir et al., 2020). However, when the gut microbiota is restored using probiotics or faecal matter transplant (FMT), the levels of IL-22 were restored to levels similar to chickens that had not undergone depletion of their gut microbiota (Yitbarek et al., 2018b). Thus, demonstrating the key part that the gut microbiota plays in viral pathogenesis. We have previously shown reduced colon microbiota alpha diversity in the acute period of AIV infection (day 2–3) in both Rhode Island Red and VALO chicken lines which did not reach the same level as in uninfected chickens by day 10 post infection, which may further influence chicken health following recovery from infection (Chrzastek et al., 2021).

Compared to the gut microbiota, the respiratory microbiota has been less well studied. It has been shown that the composition of the respiratory microbiota in chickens is also altered by host-related and environmental factors, such as age, temperature and farm location, but little is known about the relationship between the respiratory microbiota and viral infection in chickens (Glendinning et al., 2017, Wang et al., 2020, Ngunjiri et al., 2019). LPAIVs have been shown to have a tropism for respiratory tract and replicate there extensively (Guan et al., 2013, Richard et al., 2020, Abaidullah et al., 2019). Therefore, it is important to understand how the microbiota in the respiratory tract of chickens is altered following influenza A virus infection, as this may have an impact on the innate immune responses elicited there and the resulting susceptibility to infection. Understanding these changes in the respiratory microbiota could provide information useful for the development of disease prevention and treatment strategies in the future.

In this study, Rhode Island Red (RIR) chickens were used to assess temporal changes in the chicken upper respiratory track (URT) microbiota following H9N2 AIV infection. Additionally, the study aimed to assess how the healthy chicken respiratory microbiota changes during maturation up to 28 days of age.

## Materials and Methods

### Experiment design

In this study, two *in vivo* animal experiments were performed. In Experiment 1, we used 35 specific pathogen free (SPF) Rhode Island Red (RIR) chickens to assess the effect of H9N2 AIV infection on upper respiratory tract (URT) microbiota.

These birds were reared together until two weeks of age, when they were randomly separated into two groups; the control (n = 20) and H9N2 AIV infected groups (n = 15). At three weeks of age, the infected birds received 100 µl of 10^5^ PFU/ml recombinant H9N2 A/chicken/Pakistan/UDL01/01 via the intranasal route (50 µl in each nostril). On the day of challenge, 5 birds were randomly selected from the control group and culled to establish starting microbiota profile. On days 2, 4 and 10 post-challenge (p-ch), trachea samples were taken from 5 birds from both the control and infected groups. In Experiment 2, 20 SPF RIR chickens were kept in a cage together and 5 birds were randomly selected and culled at days 7, 14, 21 and 28 of age to assess URT microbiota changes over time. This data was then compared with URT microbiota results obtained from Experiment 1 to further understand the changes that occurred during infection. RIR chickens were provided as day old chicks from the National Avian Resource Facility (NARF) located at The Roslin Institute, Edinburgh, UK. The feed was provided *ad libitum* according to manufacture instruction for the chicken age. All chicks move from starter feed to grower at 3 weeks old. Control groups were housed in raised floor pens whilst AIV challenged chickens were housed in self-contained BioFlex® B50 Rigid Body Poultry isolators (Bell Isolation Systems) maintained at negative pressure. All birds were swabbed daily from day of challenge until 8 days post infection to determine viral shedding. Swabbing was carried out with sterile polyester tipped swabs (Fisher Scientific, UK) which were transferred into viral transport media (WHO, 2006), vortexed briefly, clarified by centrifugation and stored at −D80 °C prior to virus detection. The URT samples, included cranial larynx from proximal part, distally to syrinx, up to part trachea divides into bronchi. Each trachea was then cut out from larynx by sterile scalpel. Proximal part of trachea samples was used for DNA/RNA extraction and snap-frozen, whereas distal trachea was used for histology examination and thus fixed in 4% neutral buffered formaldehyde. All animal work was approved and regulated by the UK government Home Office under the project license (P68D44CF4) and reviewed by the Pirbright Animal Welfare and Ethics Review Board (AWERB). All personnel involved in the procedures were licensed by the UK Home Office. All procedures were performed in accordance with these guidelines and the study is reported in line with the ARRIVE guidelines.

### Virus propagation

Recombinant A/chicken/Pakistan/UDL01/08 H9N2 virus was generated using reverse genetics as previously described (Hoffman et al, 2000). Virus stocks were produced via passage in 10 day old embryonated chicken eggs; the allantoic fluid harvested after 48 h and titrated by plaque assay on MDCK cells (ATCC). Madin-Darby Canine Kidney (MDCK) cells (ATCC) were maintained in DMEM (Gibco-Invitrogen, Inc.) supplemented with 10% foetal bovine serum (Biosera, Inc.), 1% penicillin/streptomycin (Sigma-Aldrich, Inc.) and 1% non-essential aa (Sigma-Aldrich, Inc.).

### DNA Extraction and 16S rRNA Gene Amplification

Proximal part of trachea samples was cut and disrupted in the TissueLyser II (QIAGEN) to homogenise the tissue. Each sample was then transferred to the PowerBead tubes provided in the DNeasy PowerSoil Kit (QIAGEN). These tubes were placed in the TissueLyser LT (QIAGEN) for next 10 mins at 40 1/s. From this point onwards, the manufacturer’s instruction for the extraction of microbial genomic DNA by the DNeasy PowerSoil Kit was followed. Controls for DNA extraction reagents (negative control) and *E. coli* DH5α (TermoFisher) (positive control) were included. DNA from both controls followed the same DNA extraction protocol as described above. Controls were included to assess the level of possible contaminations and robustness of the method. The V2-V3 region of the 16S rRNA gene was amplified via a two-step nested PCR, using the protocol described by Glendinning et al. (Glendinning et al., 2017). This involved a V1-V4 region amplification using the primers 28F [5’-GAGTTTGATCNTGGCTCAG-3’] and 805R [5’-GACTACCAGGGTATCTAATC-3’]. This was followed by a short-cycle V2-V3 amplification using the primers 104F [5’-GGCGVACGGGTGAGTAA-3’] and 519R [5’- GTNTTACNGCGGCKGCTG-3’] with unique barcodes and Illumina adaptor sequences to prepare the gene segments for sequencing.

### RNA extraction and qRT-PCR

The supernatant from homogenised trachea tissue obtained at days, 2 (n=5) and 4 (n=3) post-challenge was used for RNA extraction, using the QIAmp viral RNA minikit (Qiagen) according to manufacturer instructions. For H9N2 influenza virus detection in trachea, quantitative analyses of matrix (M) were performed with primers, IAV_F (5’-AGA TGA GTC TTC TAA CCG AGG TCG-3’) and IAV_R (5’-TGC AAA AAC ATC TTC AAG TCT CTG-3’) with IAV_probe (FAM-5’ TCA GGC CCC CTC AAA GCC GA -TAMRA-3’). qRT-PCR analysis was completed using the Superscript III Platinum one-step qRT-PCR kit (Life Technologies). T7 RNA polymerase-derived transcripts from UDL-01 segment 7 were used for preparation of the standard curve. Cycling conditions were: (1) 5-min at 50 °C, (2) a 2-min step at 95 °C, and (3) 40 cycles of 3 s at 95 °C, 30 s of annealing and extension at 60 °C. Cycle threshold (CT) values were obtained using 7500 software v2.3.

### Histology

Trachea samples from control and H9N2 infected birds were taken at days 2, 4 and 10 post- challenge. This samples were fixed in 4% neutral buffered formaldehyde. In total, 21 samples were commercially prepared by ProPath® UK Limited (Hareford, UK). This included 15 samples from H9N2 infected group, five at each time point tested, accompany by control samples, 2 at each time point tested. The tissues from 21 chickens were trimmed into cassettes as per Propath’s block list and Propath’s SOPs. The tissues were processed on a Tissue Tec VIP processor, dehydrated through a series of graded alcohols, cleared in toluene and infiltrated with poly wax. The processed tissues were then embedded in poly wax and sections cut at a nominal 4µm, tissue sections were then placed on microscope glass slides. Slides were baked overnight in a 37DC oven prior to staining. Slides from each of the 21 chicken trachea samples were dewaxed in xylene, hydrated through a series of graded alcohols, and rinsed in running tap water. The slides were then stained in Mayer’s haematoxylin and washed in running tap water. The slides were then counter stained with aqueous eosin and washed in two changes of running tap water. Finally, the slides were dehydrated through a series of graded alcohols, cleared in xylene, and mounted with Pertex®. All stained slides from each of the 21 chickens were macroscopically and microscopically quality checked by ProPath® UK Limited (Hareford, UK) for the following acceptance criteria: satisfactory staining and cover slipping; presence of correct tissue/tissue area on slide, absence of section artefact, e.g. scores, creases, chatter; the presence of lesions and abnormalities noted at necropsy; consistency of embedding.

Delafield’s haematoxylin and eosin stained slides were then analysed using a light microscope. The percentage of the inner epithelial layer that had either been degraded or had the cilia on its surface stripped away was estimated independently by two people and the mean percentage damage of the two estimates was used in the statistical analysis. Two-tailed t-tests were used to assess significant differences between the control and infected groups at each time point *p < 0.05.

### 16S rRNA sequencing and Data Analysis

Libraries were analysed on a High Sensitivity DNA Chip on the Bioanalyzer (Agilent Technologies) and Qubit dsDNA HS assay (Invitrogen) and then loaded on the flow cell of the 500 cycle MiSeq Reagent Kit v2 (Illumina, USA) and pair-end sequencing (2×250 bp). Quantitative Insights into Microbial Ecology (QIIME) platform version qiime2-2019.10 was used to analyse sequencing dataset. Low-quality sequencing reads were quality trimmed and denoise using DADA2 (trim position 15 for both forward and reverse sequencing reds generated). Potential chimeric sequences were removed using UCHIME, and the remaining reads assigned to 16S rRNA operational taxonomic units (OTUs) based on 97% nucleotide similarity with the UCLUST algorithm and then classified taxonomically using the SILVA reference database (silva132-99-nb-classifer). Taxonomy was then collapsed to the genus-level. The microbial community structure was estimated by microbial biodiversity (i.e., species richness and between-sample diversity). Shannon index, phylogenetic diversity, and the observed number of species were used to evaluate alpha diversity, and the unweighted UniFrac distances were used to evaluate beta diversity. All these indices (alpha and beta diversity) were calculated by the QIIME pipeline.

### Statistical analysis of sequencing data

Kruskal Wallis pairwise statistical testing was used to assess the differences in alpha diversity in terms of the observed number of operational taxonomic units (OTUs), Faith’s phylogenetic diversity and Shannon diversity. To assess beta diversity, analysis of variance (PERMANOVA) was performed using unweighted UniFrac distances. Principal-coordinate analysis (PCoA) graphs were constructed to visualize similarity between the samples. The Linear Discriminant Analysis Effect Size (LEfSe) algorithm was used to identify differentially abundant taxa between the groups at genus level. For LEfSe analysis, depends on the experiments, different groups were assigned as comparison classes and were analysed by days. Briefly, in experiment 1, RIR control and RIR AIV infected groups were assigned as comparison classes and assessed at day 0, day 2, day 4 and day 10 post-challenge. In experiment 2, control groups were assigned as comparison classes at days 7, 14, 21, 28 and analysed all against all. LEfSe identified features that were statistically different between assigned groups and then compared the features using the non-parametric factorial Kruskal– Wallis sum-rank test (alpha value of 0.05) and Linear Discriminant Analysis (LDA) cut off value of 4.0.

## Results

### The median number of operational taxonomic units (OTUs) obtained from 16S rRNA amplicon sequencing

All samples were normalised to 62,000 sequences per sample as they had all reached plateau at this value, with exception to negative control having the total number of reads generated being only 12,670 (Suppl. Fig. 1). In Experiment 1, 4,235,519 total OTUs were assigned to control group, with a median of 232,798 OTUs and 2,909,088 OTUs to the H9N2 infected group with a median of 186,513 OTUs per sample. In Experiment 2, we obtained 3,336,323 OTUs with a median of 170,859 OTUs per sample. Due to technical issue that occurred in the sequencing stage, two samples had to be removed from analysis (one from the Experiment 1 controls at day 0 pre-challenge and one from Experiment 2, control at 28 days of age). Therefore, these groups had a sample size of four each in further analysis.

A 98 % of reads obtained from the positive control (*E. coli* DH5α) were assigned to Enterobacteriales taxa (27,3313 out of 27,9747), followed by 1.7% of reads that were assigned to Bacilli. We also found Pasteuralles, Betaproteobacteriales, Oligoflexales, Caulobacterales in our positive sample. However, for each of these taxa, no more than 200 sequencing reads were assigned (which represent less than 0.07% of total reads obtained) which might be considered an Illumina error rate rather than contamination. Two taxa, Enterobacteriales and Bacilli, was found in the negative sample. Direct comparison between negative control and experimental samples (obtained in Exp.1 and Exp 2) showed that experimental samples had at least 15x to even 105x higher number of reads assigned to Bacilli as compared to negative control, and all experimental samples were placed above contamination threshold (Suppl. Fig. 2).

### Alpha diversity indices increased during maturation in the healthy URT microbiota

We assessed the effect of maturation on alpha diversity in the URT microbiota on a weekly interval. The temporal changes were measured by number of OTUs, Faith’s phylogenetic diversity and Shannon index. Alpha diversity indices grouped by the age of the bird are shown in Figure 1. A significant increase of alpha diversity indices was seen as the chickens age, especially between 7-day old chicks and 3 weeks old birds (Fig. 1). Kruskal Wallis statistical testing showed a statistically significant difference in the number of OTUs between samples taken at day 7 and those taken at days 14, 21, and 28 (*p* < 0.01). In addition, there was a difference in the number of OTUs observed between samples taken at day 14 and day 28 (*p* < 0.05). Changes in Faith’s phylogenetic diversity were seen between day 7 and day 21, day 14 and day 28, and day 21 and day 28 (*p* < 0.05). Shannon index, significantly increased between day 7 and days 14, 21 and 28 suggesting that once the evenness was established at two-weeks of age it did not change significantly in next weeks (Suppl. Tab. 1).

Simple linear regression was also used to assess the association between alpha diversity indices and time (Suppl. Tab. 2.). The number of OTUs, Faith’s phylogenetic diversity and Shannon diversity were all significantly correlated with time (*p* < 0.001).

### A significant decrease of alpha-diversity in chicken URT microbiota was seen at days 2 and 10 post H9N2 avian influenza challenge

The effect of H9N2 AIV infection on alpha diversity in the URT microbiota was assessed at day 2, day 4 and day 10 post challenge (Figure 2). At days 2 and 10 post-challenge, all three alpha diversity metrices tested (the number of OTUs, Faith’s phylogenetic diversity and Shannon diversity) decreased in the H9N2 AIV infected group as compared to the control group with *p* < 0.001 for all three alpha indices at day 2 post-challenge and *p* < 0.05 at day 10 post-challenge (Suppl. Table 3)Faith’s phylogenetic diversity was the only metric where the H9N2 infected group was significantly reduced compared to the control group at every time point tested (Days 2, 4 and 10). The results of the Kruskal Wallis pairwise testing is shown in Suppl. Tab. 3.

### Beta diversity changes are associated with H9N2 AIV infection

To show how beta diversity changes between groups at different time points, we performed a principal coordinate analysis (PCoA) using unweighted UniFrac distances data of taxonomic composition that includes phylogenetic diversity metrics (Fig. 3). PCoA plots indicate a significant separation between control and H9N2 infected chickens at all time points tested (D2, D4, and D10 post-ch). Analysis of variance (PERMANOVA) for measuring beta-diversity showed that the H9N2 RIR infected group had significantly lower diversity as compared to control group at all time points tested (*p* < 0.05 Suppl. Tab. 4).

PCoA was also applied to assess the changes in healthy chicken URT microbiota during maturation. A significant separation in the control groups was observed over the time of birds’ maturity (Suppl. Fig. 3). Analysis of variance have shown significant differences between day 7 and days 14 (*p* < 0.01), 21 (*p* < 0.05) and 32 (*p* < 0.01). In addition, significant differences were found between day 14 and day 28 (*p* < 0.01), and day 21 and day 28 (*p* < 0.05) (Suppl. Tab. 5). Day 14 and day 21 were the only time points that was not significantly different from one another.

### Bacterial taxa that dominate the healthy URT microbiota during maturation

The relative abundance of bacterial taxa present at the phylum, class, order and family level was assessed to understand changes in healthy URT microbiota during maturation. The mean relative abundances of the dominant bacterial taxa that comprise the healthy respiratory microbiota from days 7 to 28 of age is shown on Suppl. Fig. 4. The URT microbiota is largely characterised by Firmicutes and Proteobacteria at the phylum level, and Bacilli, Clostridia, Gammaproteobacteria and Actinobacteria at the class level (Suppl. Fig. 4). Lefse analysis indicated differences in the phylogenetic distributions of the microbiota of control chickens at the different time points at OTU level (1 week, 2 weeks, 3 weeks, and 4 weeks of age) (Fig. 4). The results showed that different bacteria taxa may play a role of chicken URT maturation as we noticed that some taxa are more abundant than the other depending on chicken age. The LDA scores (LDA score [log 10]D>D4) indicated that the relative abundances of Lactobacillaceae (Lactobacillales) is the most differentially abundant in young, 7 days old chicks as compared to all other time points tested. At two weeks of age, the prevalence of Enterobacteriaceae was enriched whereas at three weeks of age, the most differentially abundant bacterial taxon was characterized by a preponderance of Bacillales (Staphylococcae), Lachnospiraceae, Gammaproteobacteria (Pseudomonales) and Actinobacteria. Clostridiales (Peptostreptococceae, Lechnospiraceae) and Lactobacillales were much more enriched at four weeks of age as compared to any other time tested. In addition, we performed two-tailed t-test analysis of changes at the phylum, class, order and family level between each of the time point tested (Suppl. Tab. 6). At the order level, there was a significantly higher relative abundance of Lactobacillales when comparing samples from day 7 to day 14, day 14 to day 21, and day 21 to day 28 (*p* < 0.01). We also noticed significant increase in Clostridiales family when comparing samples from day 7 to day 28 (*p* < 0.05) and day 14 to day 28 (*p* < 0.01).

### Lactobacillaceae (*Lactobacillus*) was the most differentially abundant taxa in URT microbiota following H9N2 AIV infection

We aimed to assess whether infection with H9N2 AIV causes the composition of URT microbiota to be dysregulated and whether certain bacterial taxa are associated with the acute or recovery phase of infection. The relative abundance of bacterial taxa present at the phylum, class, order and family level in control and infected chickens are shown in Suppl. Fig. 5. Lefse analysis indicated differences in the bacterial groups identified in the microbiota of H9N2 infected and control chickens at different time points tested (Fig. 5). The LDA scores indicated that the relative abundances of Bacilli (*Lactobacillus*) were much more enriched in H9N2 infected birds versus control at day 2 post-challenge (Fig. 5A) (LDA score [log 10]>4). Additionally, two-tailed t-test analysis of changes at the phylum level, showed significant growth in the relative proportion of Firmicutes at day 2 post-ch in the infected group compared to the control group due to the significant increase in the relative abundance of the family Lactobacillaceae (as well as its class Bacilli and order Lactobacillales), as this family highly dominates the respiratory microbiota at this time point (Suppl. Tab. 7). The most differentially abundant bacterial taxon in control birds was characterized by a preponderance of Staphylococcaceae (*Staphylococcus*) and Actinobacteria (LDA score [log10]>4) at day 2 post-challenge.

Furthermore, we observed differential abundance of bacterial taxa between H9N2 infected and control birds at day 10 post-ch. The LDA scores indicated that the relative abundances of Lactobacillaceae (Lactobacillales) and Enterobacteriaceae (Enterobacteriales) were much more enriched in H9N2 infected birds versus control at recovery phase of infection (Fig. 5B) and the most differentially abundant bacteria taxa (LDA score [log 10]>4.0). A significant increase in the relative abundance of the family Lactobacillaceae at day 10 post-ch was also seen by two-tailed t-test analysis (Suppl. Tab. 7). The control birds were characterized mainly by a preponderance of Bacillales (Staphylococcaceae), Actinobacteria, Gammaproteobacteria, Propionibacteriaceae, Corynebacteriales. No differentially abundant taxa were seen above LDA score [log10]>4 at day 4 post-challenge besides Enterobacteriales enriched in H9N2 infected birds (Suppl. Fig. 6).

### The influenza A virus matrix gene is detected at days 2 and 4 post-challenge in the tracheas of the H9N2 AIV chickens

We performed qRT-PCR on RNA extracted from H9N2 AIV infected trachea samples using primers specific for M-gene to verify that the virus is replicating in the respiratory tract. Copies of the M- gene were detected in all samples tested at days 2 and 4 post-challenge (Suppl. Fig 7).

### H9N2 AIV infection is associated with increased damage to the inner epithelial cell layer in tracheas at day 4 but not at day 2 and day 10 post-challenge

To assess the damage to the inner epithelial layer of the respiratory tract, samples of the tracheas taken at days 2, 4 and 10 post-ch with H9N2 AIV from both control and infected birds and were examined for inner epithelial cell layer damage. As compared to healthy inner epithelial layer of trachea section (6A), the damage appeared to occur in two ways; the degradation of the epithelial layer (Fig. 6B) or where the cilia on the cell surface was damaged, but the epithelial layer was left intact. Independent estimates were made to the percentage of the inner epithelial layer that was damaged (Fig. 6C). Two-tailed t-tests were used to assess where there were significant differences between the control and infected groups. At days 2 and 10 post-challenge, no differences were shown between the control and infected groups (p = 0.4504 and p = 0.92829, respectively). The H9N2 infected group at day 4 post-challenge showed significantly more damage to the inner epithelial layer (p = 0.048518) as compared to the control group at the same time point tested. It is, however, worth to notice that we also observed substantial damage to the tissues in the control samples collected at day 4 post-ch which was not seen at day 2 and day 10 post-ch at the control groups (Fig. 6C).

## Discussion

We have demonstrated shown that the chicken upper respiratory (URT) microbiota is altered at acute and recovery phase of AIV H9N2 infection. Both, alpha and beta diversity indices were continuously compromised following AIV infection. Furthermore, the composition of bacteria taxa also differs, suggesting that different bacteria taxa might play a role in acute and recovery phase from AIV infection. The presence of the viral M-gene in the tracheas of chickens infected with H9N2 AIV was seen during the acute, infectious phase (days 2 and 4 post-challenge) verifying the replication of influenza A virus in tracheas at these time points tested. In addition, the highest levels of oral viral shedding were also detected at day 2 post-challenge which ceased by day 6 (Chrzastek et al., 2021) providing further evidence on active infection. During the acute phase of infection, the most significant losses in species richness, evenness and phylogenetic diversity were seen, which continued to be compromised also at recovery phase of infection. In our previous study (Chrzastek et al., 2021), we have seen similar trend where all colon microbiota of H9N2 AIV infected birds lost their overall richness at the acute phase of infection, however, species richness was restored in the recovery phase through the predominant bacteria becoming more dominant within the microbiota which we did not seen in URT microbiota. Therefore, the typical levels of diversity in the upper respiratory microbiota are not quickly restored after the infectious phase.

The major bacterial families that characterise the healthy upper respiratory microbiota during maturation are Lactobacillaceae, Staphylococcaceae, Lachnospiraceae, Enterobacteriaceae and Ruminococcaceae. This is in agreements with previous studies, which have shown that members of Lactobacillales are the most abundant in the respiratory microbiota, along with others like Enterobacteriales (Sohail et al., 2015, Glendinning et al., 2017, Kawaguchi et al., 1992). It has also been shown previously that Lactobacillaceae members are early colonisers of the chicken respiratory tract but become less abundant as the microbiota becomes more diverse (Glendinning et al., 2017). This is supported in our findings, as Lactobacillaceae is the most abundant at 7 days of age, but it becomes less prominent over time as species evenness advances over time. However, the composition of bacteria in URT can changed during the H9N2 AIV infection as we have shown in this study. In the acute phase of H9N2 AIV infection, Lactobacillaceae highly dominates the upper respiratory microbiota. Furthermore, Lactobacillaceae were still predominant bacteria at recovery phase of infection suggesting on important role of this family in surviving influenza infection. Lactobacillaceae is a family of gram-positive lactic acid bacteria that has been shown to act as a probiotic (Wells, 2011). Members of the Lactobacillaceae family, particularly the genus Lactobacillus, have been shown to regulate the immune system and can activate innate and adaptive immune responses (Wells, 2011, Nouri Gharajalar et al., 2020). Interestingly, in our previous study that regards to the colon microbiota we have noticed that although Firmicutes phylum was the most differentially abundant between infected and non-infected individuals, the Lactobacillales was missing at recovery phase of infection, and this was seen in two genetically distinct chicken breeds infected with H9N2 AIV (Chrzastek et al., 2021). The exact reason why Lactobacillaceae is so enriched in the acute and recovery phase of H9N2 AIV infection in the chicken respiratory tract and missing at recovery phase in colon microbiota needs to be further investigated. However, this is the first report showing that there might be some potential interactions, cross-talk between gut-respiratory microbiota during active viral infection in chicken.

Next question we have asked in this study was, whether the reason for dysbiosis of the respiratory microbiota observed during the H9N2 AIV infection relates to physical disruption of the epithelial cells lining of the respiratory tract, which in turn could potentially causes dysbiosis of the microbial communities that reside on it. No difference in epithelial cell lining was observed between the control and infected groups at days 2 and 10 post-challenge suggesting that there must be another factor/s that corresponded to microbiota dysbiosis at this time point tested. We hypothesised that the reason for dysbiosis particularly at acute phase of infection might be connected with innate immune response elicit towards the replicating influenza A virus which could potentially directly impair the presence of some commensal bacteria or it might have created an inflammatory environment where only some type of bacteria can survive. This suggests that bacteria might be either killed directly by immune responses or bacteria that are better adapted to persist in harsher conditions are able to thrive, while other species die out. The association between damage to the inner epithelial layer and H9N2 AIV infection was seen at day 4 post-challenge, however, this do not substantially associate with differences in microbial composition between the control and infected chickens observed in this study. It is also possible that the reason why day 4 post-challenge showed less difference between the control and infected groups is because some of the trachea samples in the control group also showed damage to the inner epithelial layer (which we are not able to explain) and thus resulted in alterations to the microbiota that reduced its species richness and evenness and, therefore, making them more similar to the microbiota following challenge with H9N2 AIV. The damage to the tissues in infected samples, tho, could be connected with antiviral mechanism regulated by an interplay between different cytokine and interferon types. Morris et al. (2023) have shown upregulation of genes responsible for defence and inflammatory responses at day 5 in lungs following H9N2 challenge in chicken.

In addition to assessing changed in URT microbiota, during the AIV H9N2 infection, we have also shown that one of the factors that determines the composition of the healthy respiratory microbiota in URT is age, as we observed alpha diversity indices increasing over time and differential enrichment of bacterial taxa as chickens mature up to 28 days of age. A significant positive correlation between alpha diversity indices and increasing age was observed, with the most significant period being between day 7 and days 14, 21 and 28. Beta diversity analysis supported our finding that that the microbial diversity is shaped by ageing. Interestingly, no significant changes in beta diversity were found between days 14 and 21 of age, neither in alpha diversity indices (OTUs and phylogenetic diversity) at this time point tested, indicating that the least substantial changes occur between week two and three of age. Although our control Experiment 2 finished at birds age of 28 days, we observed significant separation between the day 0 and 10 control groups in our challenge experiment (Experiment 1). This provides further evidence for age being a factor that influences the respiratory microbiota, as it suggests that the microbial composition in the respiratory tract has significantly evolved during maturation from days 21 to 31 of age, which goes further beyond the timeframe of age we studied in chickens for Experiment 2. It was previously shown that the respiratory tract of chickens has been shown to be populated by a diverse community of commensal bacteria, whose composition can be altered by a number of host- and environmental-related factors (Glendinning et al., 2017, Kawaguchi et al., 1992, Ngunjiri et al., 2019).

In conclusion, this is the first study showing alteration of upper respiratory track microbiota flowing the H9N2 AIV infection in chickens. We observed that alpha and beta diversity in the respiratory microbiota change in both the acute and recovery phase of influenza A virus infection and that there is dysbiosis in the abundances of different bacterial taxa at different time points. Lack of homeostasis on the upper respiratory track seems to be connected with different factors and might change over the time of infection. Dysbiosis might be more associated with anti-viral response, which could then directly or indirectly (by changing the environment) affect microbiota on mucosal surfaces, and less with physical damage done to the inner epithelial layer cells by replicating virus. The family Lactobacillaceae is most strongly associated with infected chickens compared to the controls during the acute phase and continues to be major player in the respiratory microbiota into the recovery phase, while the abundance of less dominant families is reduced to levels lower than in control chickens. Although we do not know why Lactobacillaceae family was the main player during the AIV infection in the upper respiratory track of chickens, as this needs to be investigated further, this study might suggest that supplementation with Lactobacillaceae could potentially aid recovery from H9N2 influenza infection.

---

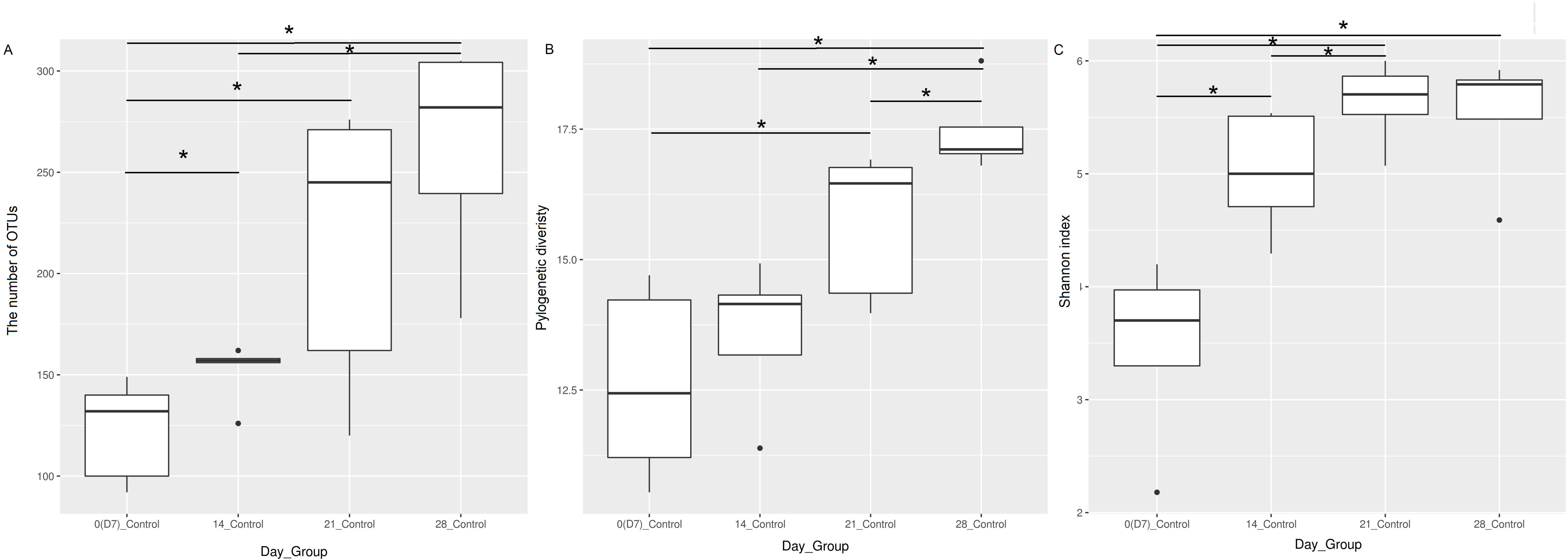

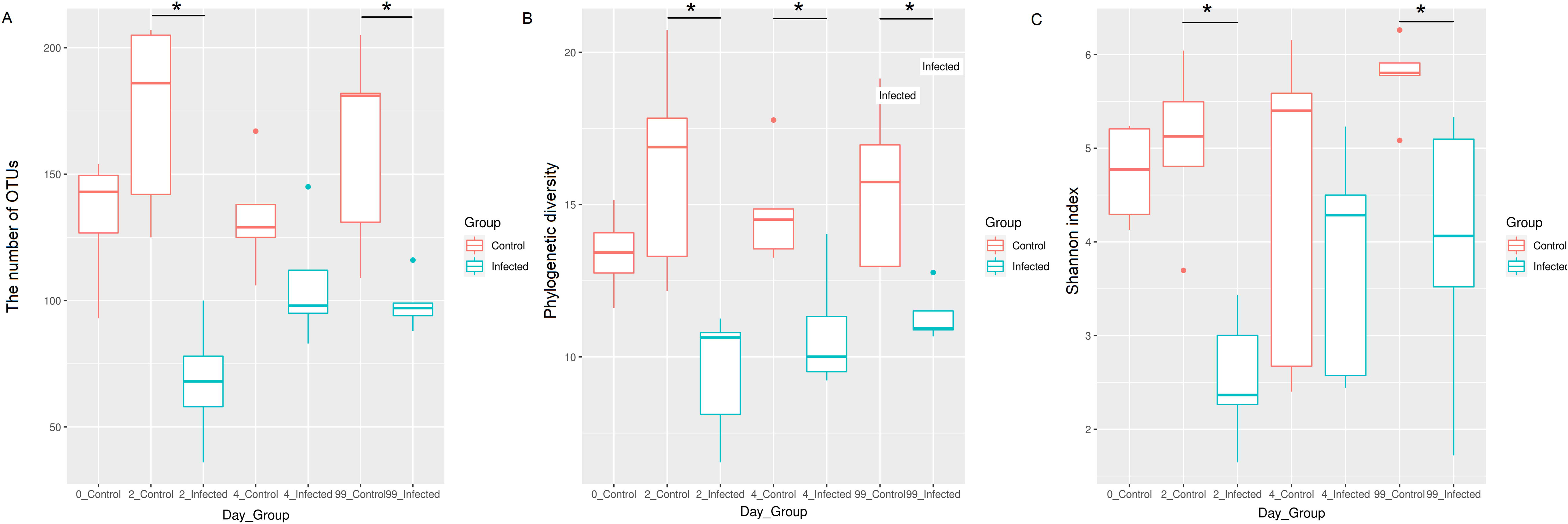

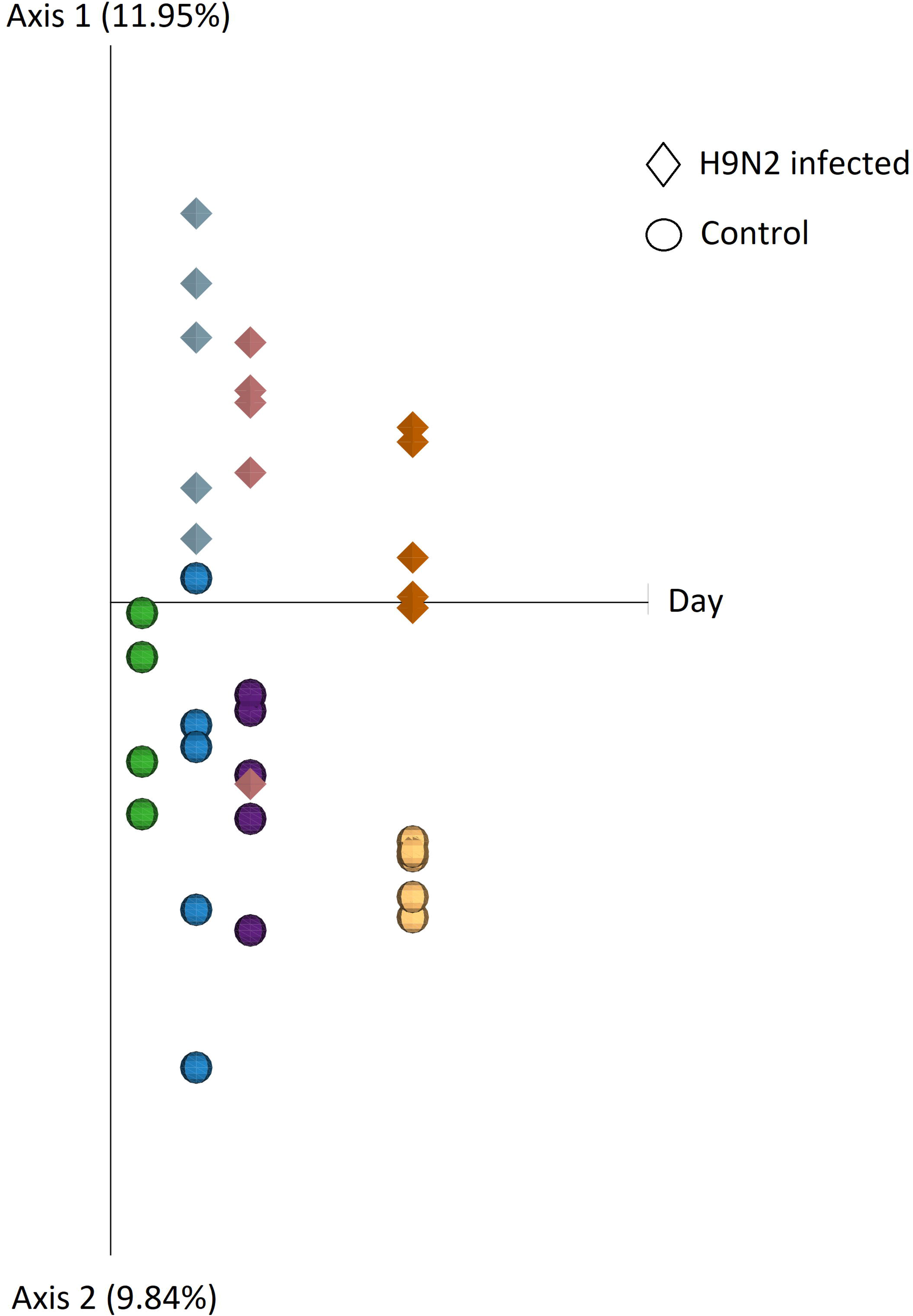

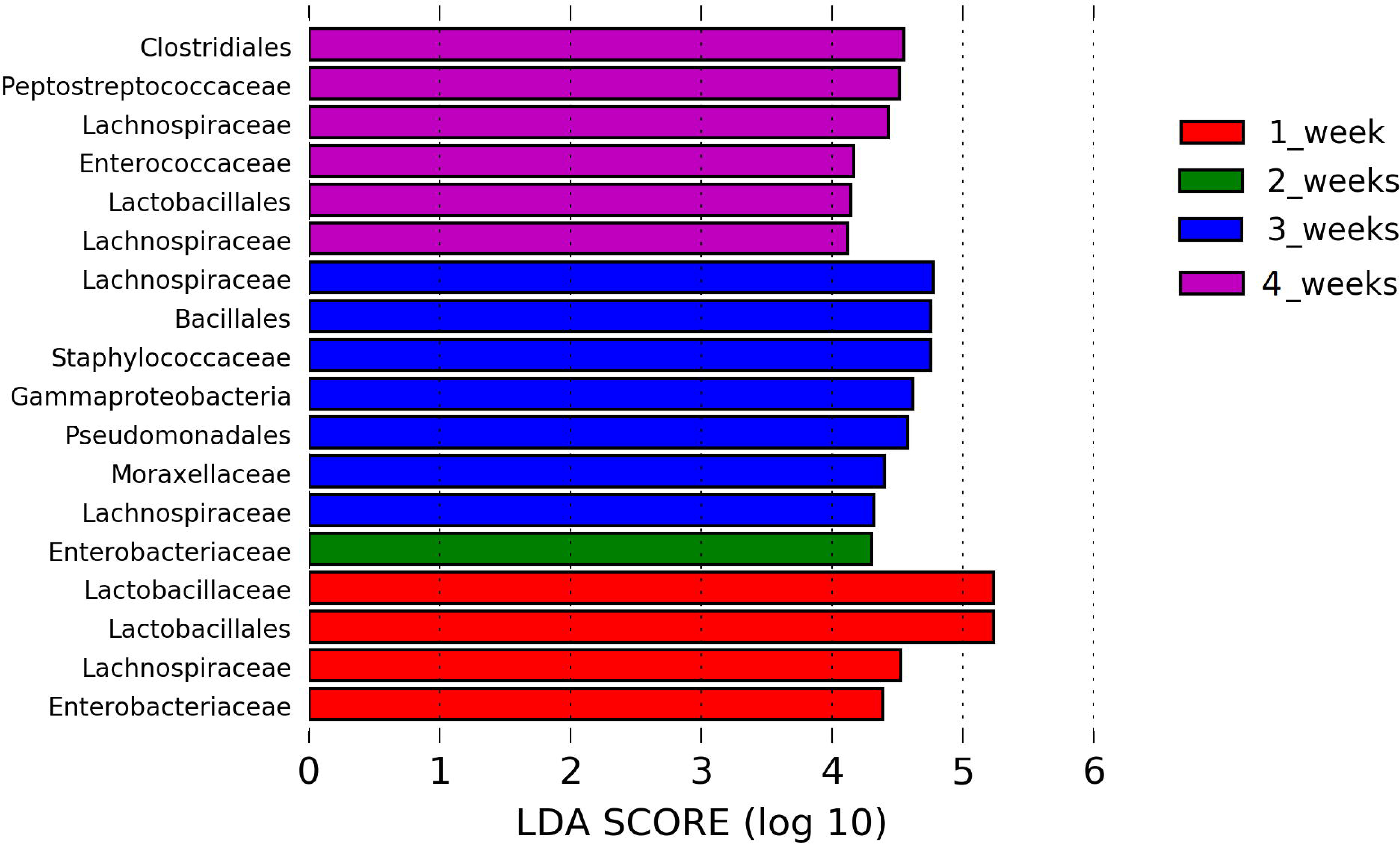

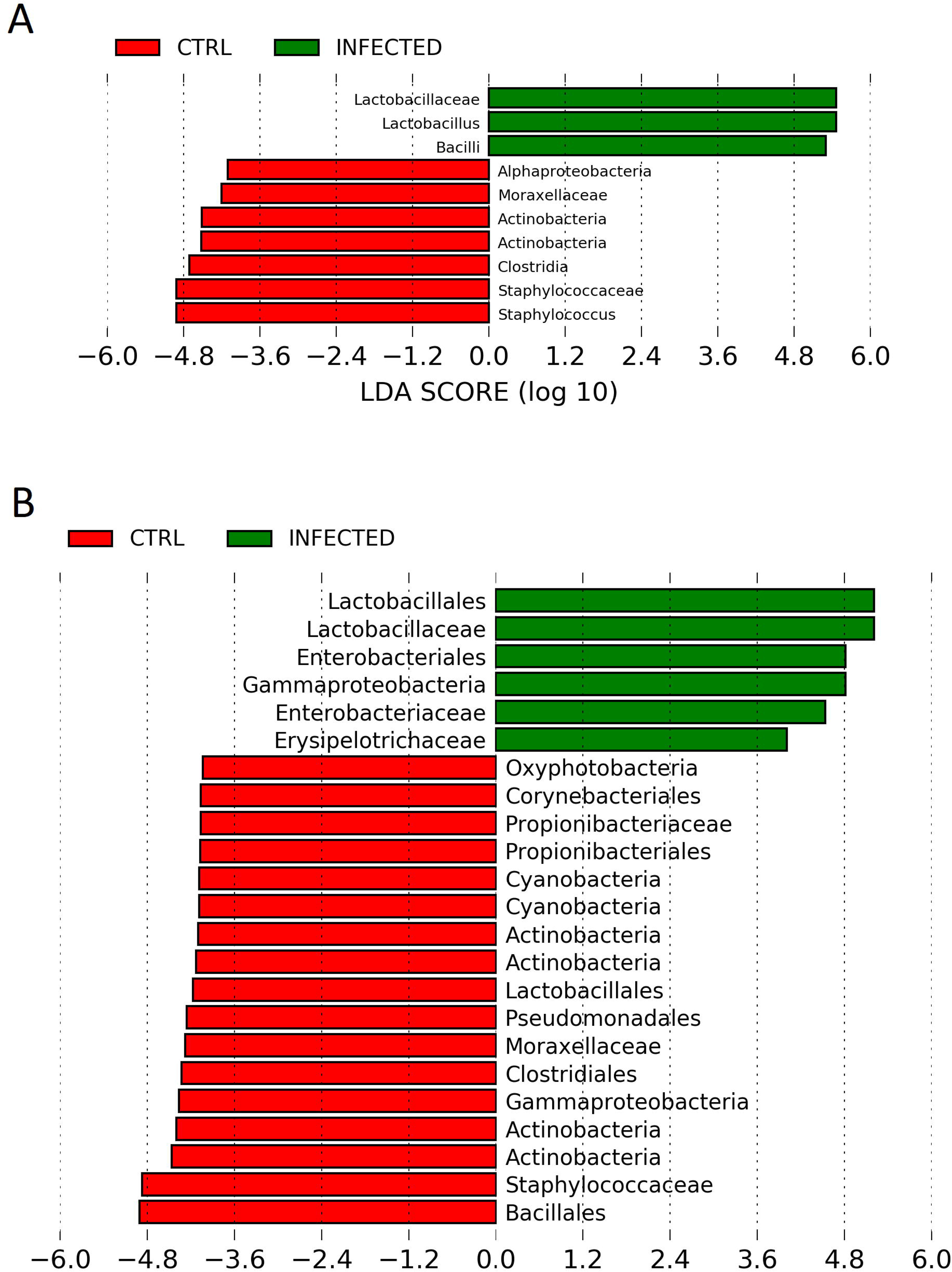

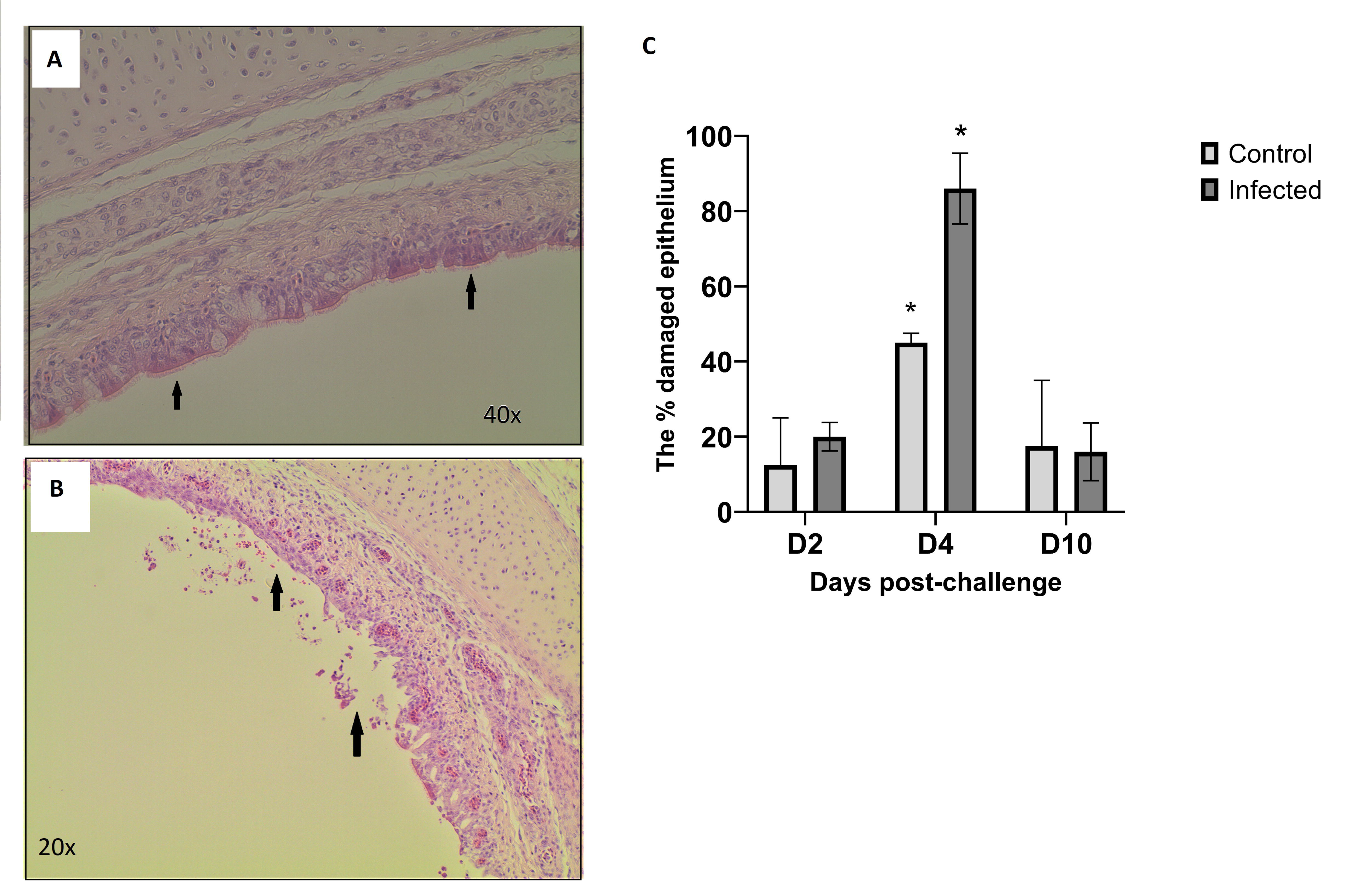

## References

Abaidullah, M., Peng, S., Kamran, M., Song, X. & Yin, Z. 2019. Current Findings on Gut Microbiota Mediated Immune Modulation against Viral Diseases in Chicken. Viruses, 11.

Alexander, D. J. 2007. An overview of the epidemiology of avian influenza. Vaccine, 25, 5637–44.

Bavananthasivam, J., Astill, J., Matsuyama-Kato, A., Taha-Abdelaziz, K., Shojadoost, B. & Sharif, S. 2021. Gut microbiota is associated with protection against Marek’s disease virus infection in chickens. Virology, 553, 122–130.

Bottcher-Friebertshauser, E., Klenk, H. D. & Garten, W. 2013. Activation of influenza viruses by proteases from host cells and bacteria in the human airway epithelium. Pathog Dis, 69, 87–100.

Chrzastek, K., Leng, J., Zakaria, M. K., Bialy, D., LA Ragione, R. & Shelton, H. 2021. Low pathogenic avian influenza virus infection retards colon microbiota diversification in two different chicken lines. Anim Microbiome, 3, 64.

Clavijo, V. & Florez, M. J. V. 2018. The gastrointestinal microbiome and its association with the control of pathogens in broiler chicken production: A review. Poult Sci, 97, 1006–1021.

Glendinning, L., Mclachlan, G. & Vervelde, L. 2017. Age-related differences in the respiratory microbiota of chickens. PLoS One, 12, e0188455.

Gu, M., Xu, L., Wang, X. & Liu, X. 2017. Current situation of H9N2 subtype avian influenza in China. Vet Res, 48, 49.

Guan, J., Fu, Q., Chan, M. & Spencer, J. L. 2013. Aerosol transmission of an avian influenza H9N2 virus with a tropism for the respiratory tract of chickens. Avian Dis, 57, 645–9.

Harmon, K., Bower, L., Kim, W. I., Pentella, M. & Yoon, K. J. 2010. A matrix gene-based multiplex real-time RT-PCR for detection and differentiation of 2009 pandemic H1N1 and other influenza A viruses in North America. Influenza Other Respir Viruses, 4, 405–10.

Hoffmann, E., Neumann, G., Hobom, G., Webster, R. G. & Kawaoka, Y. 2000. "Ambisense" approach for the generation of influenza A virus: vRNA and mRNA synthesis from one template. Virology, 267, 310–7.

Kawaguchi, I., Hayashidani, H., Kaneko, K., Ogawa, M. & Benno, Y. 1992. Bacterial flora of the respiratory tracts in chickens with a particular reference to Lactobacillus species. J Vet Med Sci, 54, 261–7.

Keir, M., Yi, Y., Lu, T. & Ghilardi, N. 2020. The role of IL-22 in intestinal health and disease. J Exp Med, 217, e20192195.

Kers, J. G., Velkers, F. C., Fischer, E. A. J., Hermes, G. D. A., Stegeman, J. A. & Smidt, H. 2018. Host and Environmental Factors Affecting the Intestinal Microbiota in Chickens. Front Microbiol, 9, 235.

Lee, D. H., Swayne, D. E., Sharma, P., Rehmani, S. F., Wajid, A., Suarez, D. L. & Afonso, C. 2016. H9N2 low pathogenic avian influenza in Pakistan (2012-2015). Vet Rec Open, 3, e000171.

Li, C., Wang, S., Bing, G., Carter, R. A., Wang, Z., Wang, J., Wang, C., Wang, L., Wu, G., Webster, R. G., Wang, Y., Sun, H., Sun, Y., Liu, J. & Pu, J. 2017. Genetic evolution of influenza H9N2 viruses isolated from various hosts in China from 1994 to 2013. Emerg Microbes Infect, 6, e106.

Li, H., Liu, X., Chen, F., Zuo, K., Wu, C., Yan, Y., Chen, W., Lin, W. & Xie, Q. 2018. Avian Influenza Virus Subtype H9N2 Affects Intestinal Microbiota, Barrier Structure Injury, and Inflammatory Intestinal Disease in the Chicken Ileum. Viruses, 10.

Morris, K. M., Mishra, A., Raut, A. A., Gaunt, E. R., Borowska, D., Kuo, R. I., Wang, B., Vijayakumar, P., Chingtham, S., Dutta, R., Baillie, K., Digard, P., Vervelde, L., Burt, D. W. & Smith, J. 2023. The molecular basis of differential host responses to avian influenza viruses in avian species with differing susceptibility. Front Cell Infect Microbiol, 13, 1067993.

Mostafa, A., Abdelwhab, E. M., Mettenleiter, T. C. & Pleschka, S. 2018. Zoonotic Potential of Influenza A Viruses: A Comprehensive Overview. Viruses, 10.

Ngunjiri, J. M., Taylor, K. J. M., Abundo, M. C., Jang, H., Elaish, M., Kc, M., Ghorbani, A., Wijeratne, S., Weber, B. P., Johnson, T. J. & Lee, C. W. 2019. Farm Stage, Bird Age, and Body Site Dominantly Affect the Quantity, Taxonomic Composition, and Dynamics of Respiratory and Gut Microbiota of Commercial Layer Chickens. Appl Environ Microbiol, 85.

Nouri Gharajalar, S., Mirzai, P., Nofouzi, K. & Madadi, M. S. 2020. Immune enhancing effects of Lactobacillus acidophilus on Newcastle disease vaccination in chickens. Comp Immunol Microbiol Infect Dis, 72, 101520.

Peacock, T. H. P., James, J., Sealy, J. E. & Iqbal, M. 2019. A Global Perspective on H9N2 Avian Influenza Virus. Viruses, 11.

Richard, M., Van Den Brand, J. M. A., Bestebroer, T. M., Lexmond, P., De Meulder, D., Fouchier, R. A. M., Lowen, A. C. & Herfst, S. 2020. Influenza A viruses are transmitted via the air from the nasal respiratory epithelium of ferrets. Nat Commun, 11, 766.

Sohail, M. U., Hume, M. E., Byrd, J. A., Nisbet, D. J., Shabbir, M. Z., Ijaz, A. & Rehman, H. 2015. Molecular analysis of the caecal and tracheal microbiome of heat-stressed broilers supplemented with prebiotic and probiotic. Avian Pathol, 44, 67–74.

Spackman, E. 2020. A Brief Introduction to Avian Influenza Virus. Methods Mol Biol, 2123, 83–92.

Sun, Y. & Liu, J. 2015. H9N2 influenza virus in China: a cause of concern. Protein Cell, 6, 18–25.

Wang, M., Lin, X., Jiao, H., Uyanga, V., Zhao, J., Wang, X., Li, H., Zhou, Y., Sun, S. & Lin, H. 2020. Mild heat stress changes the microbiota diversity in the respiratory tract and the cecum of layer-type pullets. Poult Sci, 99, 7015–7026.

Wells, J. M. 2011. Immunomodulatory mechanisms of lactobacilli. Microb Cell Fact, 10 Suppl 1, S17.

Yitbarek, A., Alkie, T., Taha-Abdelaziz, K., Astill, J., Rodriguez- LeCOmpte, J. C., Parkinson, J., Nagy, E. & Sharif, S. 2018. Gut microbiota modulates type I interferon and antibody-mediated immune responses in chickens infected with influenza virus subtype H9N2. Benef Microbes, 9, 417–427.

yitbarek, A., taha-abdelaziz, K., hodgins, D. C., read, L., nagy, E., weese, J. S., caswell, J. L., parkinson, J. & sharif, S. 2018. Gut microbiota-mediated protection against influenza virus subtype H9N2 in chickens is associated with modulation of the innate responses. Sci Rep, 8, 13189.

yitbarek, A., weese, J. S., alkie, T. N., parkinson, J. & sharif, S. 2018. Influenza A virus subtype H9N2 infection disrupts the composition of intestinal microbiota of chickens. FEMS Microbiol Ecol, 94.

WHO, 2006

