## Supplementary Tables for "Alteration of the chicken upper respiratory microbiota, following H9N2 avian influenza virus infection": Suppl. Tables_Microbiota_ChrzastekK.docx

**S1. Kruskal Wallis pairwise testing of the association between alpha diversity and age in the healthy respiratory microbiome in chickens.** Kruskal Wallis pairwise testing was used to assess the changes in alpha diversity indices between time points in Experiment 2. Samples in Experiment 2 were taken at 7, 14, 21 and 28 days of age from healthy RIR chickens. The alpha diversity in the respiratory microbiome was tested using the number of OTUs detected, Faith’s phylogenetic diversity and Shannon diversity.

| **Alpha Diversity** | **Group 1** | **Group 2** | **H** | **p-value** | **q-value** |
| --- | --- | --- | --- | --- | --- |
| The number of OTUs | Day 7 | Day 14 | 3.93818182 | 0.04720177 | 0.11981987 |
|  | Day 7 | Day 21 | 3.93818182 | 0.04720177 | 0.11981987 |
|  | Day 7 | Day 28 | 6 | 0.01430588 | 0.05974321 |
|  | Day 14 | Day 21 | 2.15121951 | 0.1424567 | 0.21984893 |
|  | Day 14 | Day 28 | 6 | 0.01430588 | 0.05974321 |
|  | Day 21 | Day 28 | 1.5 | 0.22067136 | 0.29723081 |
| Faith’s Phylogenetic Diversity | Day 7 | Day 14 | 0.88363636 | 0.34720764 | 0.44932753 |
|  | Day 7 | Day 21 | 3.93818182 | 0.04720177 | 0.12461267 |
|  | Day 7 | Day 28 | 6 | 0.01430588 | 0.06721111 |
|  | Day 14 | Day 21 | 3.15272727 | 0.07580017 | 0.16676038 |
|  | Day 14 | Day 28 | 6 | 0.01430588 | 0.06721111 |
|  | Day 21 | Day 28 | 4.86 | 0.02748634 | 0.07777034 |
| Shannon Diversity | Day 7 | Day 14 | 6.81818182 | 0.00902344 | 0.08507814 |
|  | Day 7 | Day 21 | 6.81818182 | 0.00902344 | 0.08507814 |
|  | Day 7 | Day 28 | 6 | 0.01430588 | 0.08961482 |
|  | Day 14 | Day 21 | 3.93818182 | 0.04720177 | 0.15013056 |
|  | Day 14 | Day 28 | 2.16 | 0.14164469 | 0.26259733 |
|  | Day 21 | Day 28 | 0 | 1 | 1 |

**S2. Simple linear regression of the association between alpha diversity in the respiratory microbiome and the age of chickens.** Alpha diversity metrices (the number of OTUs, Faith’s phylogenetic diversity and Shannon Diversity) were used as the dependent variable in simple linear regression to determine whether there is correlation between the species richness in the respiratory microbiome and chicken age. Samples tested were taken at 7, 14, 21 and 28 days of age.

| **Alpha Diversity** | **R2** | **Slope (Y value)** | **Equation** | **P Value** |
| --- | --- | --- | --- | --- |
| Observed Number of OTUs | 0.5998 | 1.356 | Y = 6.844*X + 67.85 | <0.0001 |
| Faith’s Phylogenetic Diversity | 0.6538 | 0.04167 | Y = 0.2361*X + 10.70 | <0.0001 |
| Shannon Diversity | 0.5684 | 0.02132 | Y = 0.1009*X + 3.167 | 0.0002 |

**S3. Kruskal Wallis pairwise testing of the differences in alpha diversity between the control and H9N2 AIV infected respiratory microbiome in chickens.** Kruskal Wallis pairwise testing was used to assess how alpha diversity indices change between control groups and groups challenged with H9N2 AIV A/chicken/Pakistan/UDL01/08 at days 2 (D2), 4 (D4) and 10 (D10) post-challenge in RIR chickens (Experiment 1). The alpha diversity in the respiratory microbiome was tested using the number of OTUs detected, Faith’s phylogenetic diversity and Shannon diversity.

| **Alpha Diversity** | **Days Post-Challenge** | **H** | **p-value** | **q-value** |
| --- | --- | --- | --- | --- |
| The number of OTUs | D2 | 6.81818182 | 0.00902344 | 0.05974321 |
|  | D4 | 2.45454546 | 0.11718509 | 0.21483933 |
|  | D10 | 5.77090909 | 0.0162936 | 0.05974321 |
| Faith’s Phylogenetic Diversity | D2 | 6.81818182 | 0.00902344 | 0.06721111 |
|  | D4 | 4.81090909 | 0.02828012 | 0.07777034 |
|  | D10 | 6.81818182 | 0.00902344 | 0.06721111 |
| Shannon Diversity | D2 | 6.81818182 | 0.00902344 | 0.08507814 |
|  | D4 | 0.88363636 | 0.34720764 | 0.44932753 |
|  | D10 | 4.81090909 | 0.02828012 | 0.10979342 |

**S4. Analysis of variance (PERMANOVA) of beta diversity between the healthy respiratory microbiome at different ages.** To assess beta diversity between different time points in Experiment 2, unweighted UniFrac distances were used in PERMANOVA. Samples in Experiment 2 was taken at 7, 14, 21 and 28 days of age.

| **Group 1** | **Group 2** | **pseudo-F** | **p-value** | **q-value** |
| --- | --- | --- | --- | --- |
| Day 7 | Day 14 | 1.843418132 | 0.008 | 0.022578947 |
| Day 7 | Day 21 | 2.455765667 | 0.012 | 0.022578947 |
| Day 7 | Day 28 | 3.697054644 | 0.005 | 0.022578947 |
| Day 14 | Day 21 | 1.28880597 | 0.052 | 0.073021277 |
| Day 14 | Day 28 | 2.580119787 | 0.006 | 0.022578947 |
| Day 21 | Day 28 | 1.821964447 | 0.026 | 0.039906977 |

**S5. Analysis of variance (PERMANOVA) of beta diversity between control and H9N2 AIV infected respiratory microbiome.** To assess beta diversity between different time points in Experiment 1, unweighted UniFrac distances were used in PERMANOVA. Samples were taken at day 0 (D0) pre-challenge and at days 2 (D2), 4 (D4) and 10 (D10) post-challenge in control chickens and chickens challenged with H9N2 AIV A/chicken/Pakistan/UDL01/08.

| Group 1 | Group 2 | pseudo-F | p-value | q-value |
| --- | --- | --- | --- | --- |
| CTRL D0 | CTRL D2 | 1.12756 | 0.208 | 0.2288 |
| CTRL D0 | Infected D2 | 1.955375 | 0.009 | 0.022578947 |
| CTRL D0 | Infected D4 | 2.002263 | 0.028 | 0.042 |
| CTRL D0 | Infected D10 | 1.750084 | 0.006 | 0.022578947 |
| CTRL D2 | Infected D2 | 2.075569 | 0.005 | 0.022578947 |
| CTRL D2 | Infected D4 | 1.899262 | 0.03 | 0.043043478 |
| CTRL D2 | Infected D10 | 1.712934 | 0.015 | 0.025384615 |
| CTRL D4 | Infected D4 | 1.884073 | 0.026 | 0.039906977 |
| CTRL D4 | Infected D10 | 1.580944 | 0.009 | 0.022578947 |
| CTRL D10 | Infected D10 | 2.146785 | 0.012 | 0.022578947 |

**S6. Two-tailed t-test analysis of changes in relative abundances of the dominant bacterial taxa in the healthy respiratory microbiota during maturation.** Two-tailed t-tests were used to assess how the relative abundances of bacterial taxa at phylum, class, order and family level change between different ages during maturation. Samples of the respiratory microbiota were taken at day 7 (D7), 14 (D14), 21 (D21) and 28 (D28). Only bacterial taxa that accounted for more than 1% of total sequences were included in the graphs. All other bacterial taxa were categorised as ‘Other’.

| **Bacterial Taxa** | **Group Pairings** | | | | | | | | | | | |
| --- | --- | --- | --- | --- | --- | --- | --- | --- | --- | --- | --- | --- |
|  | **D7** | **D14** | **D7** | **D21** | **D7** | **D28** | **D14** | **D21** | **D14** | **D28** | **D21** | **D28** |
| **Phylum Level** | | | | | | | | | | | | |
| Actinobacteria | 0.070479 | | 0.050977 | | 0.23694 | | 0.710197 | | 0.462505 | | 0.315832 | |
| Firmicutes | 0.067229 | | 0.162042 | | 0.878503 | | 0.582231 | | 0.028109 | | 0.081141 | |
| Proteobacteria | 0.094927 | | 0.195898 | | 0.810353 | | 0.563305 | | 0.038984 | | 0.075587 | |
| Other | 0.257428 | | 0.963172 | | 0.936832 | | 0.203231 | | 0.229532 | | 0.880739 | |
| **Class Level** | | | | | | | | | | | | |
| Actinobacteria | 0.063836 | | 0.049891 | | 0.241134 | | 0.744997 | | 0.414006 | | 0.298763 | |
| Bacilli | 0.253242 | | 0.052482 | | 0.157876 | | 0.180543 | | 0.546244 | | 0.427871 | |
| Clostridia | 0.341719 | | 0.557329 | | 0.042995 | | 0.195407 | | 0.00154 | | 0.318713 | |
| Alphaproteobacteria | 0.006441 | | 0.03074 | | 0.166074 | | 0.673572 | | 0.433652 | | 0.358379 | |
| Gammaproteobacteria | 0.112765 | | 0.241603 | | 0.740325 | | 0.505886 | | 0.039521 | | 0.072502 | |
| Other | 0.857599 | | 0.817597 | | 0.432822 | | 0.995679 | | 0.557588 | | 0.454002 | |
| **Order Level** | | | | | | | | | | | | |
| Corynebacteriales | 0.189263 | | 0.153678 | | 0.373971 | | 0.728018 | | 0.321629 | | 0.315377 | |
| Propionibacteriales | 0.10657 | | 0.046058 | | 0.369735 | | 0.73526 | | 0.508449 | | 0.325371 | |
| Bacillales | 0.001022 | | 0.009621 | | 0.010299 | | 0.126793 | | 0.141482 | | 0.075967 | |
| Lactobacillales | 0.104493 | | 0.006265 | | 0.098038 | | 0.019124 | | 0.747027 | | 0.026919 | |
| Clostridiales | 0.341719 | | 0.557329 | | 0.042995 | | 0.195407 | | 0.00154 | | 0.318713 | |
| Betaproteobacteriales | 0.314155 | | 0.221674 | | 0.969566 | | 0.395285 | | 0.371286 | | 0.283636 | |
| Enterobacteriales | 0.221997 | | 0.857193 | | 0.761448 | | 0.174287 | | 0.10803 | | 0.408671 | |
| Pseudomonadales | 0.137666 | | 0.06184 | | 0.73149 | | 0.202939 | | 0.100799 | | 0.080402 | |
| Other | 0.002262 | | 0.039035 | | 0.238429 | | 0.629033 | | 0.630406 | | 0.49778 | |
| **Family Level** | | | | | | | | | | | | |
| Corynebacteriaceae | 0.189263 | | 0.158282 | | 0.373971 | | 0.722767 | | 0.321629 | | 0.322238 | |
| Propionibacteriaceae | 0.12948 | | 0.03166 | | 0.337235 | | 0.869919 | | 0.575345 | | 0.40401 | |
| Staphylococcaceae | 0.0002 | | 0.008229 | | 0.023666 | | 0.09715 | | 0.080057 | | 0.059108 | |
| Lactobacillaceae | 0.082136 | | 0.005879 | | 0.064225 | | 0.017344 | | 0.534994 | | 0.063102 | |
| Family XI | 0.356339 | | 0.983505 | | 0.456621 | | 0.357042 | | 0.376897 | | 0.375385 | |
| Lachnospiraceae | 0.079583 | | 0.37311 | | 0.056789 | | 0.081177 | | 0.001582 | | 0.704507 | |
| Ruminococcaceae | 0.196306 | | 0.467582 | | 0.759995 | | 0.108931 | | 0.005192 | | 0.354574 | |
| Burkholderiaceae | 0.311983 | | 0.223401 | | 0.86452 | | 0.399149 | | 0.20495 | | 0.240814 | |
| Enterobacteriaceae | 0.221997 | | 0.857193 | | 0.761448 | | 0.174287 | | 0.10803 | | 0.408671 | |
| Moraxellaceae | 0.270612 | | 0.066002 | | 0.654778 | | 0.194091 | | 0.193102 | | 0.078969 | |
| Pseudomonadaceae | 0.052892 | | 0.197058 | | 0.276808 | | 0.722791 | | 0.153514 | | 0.316791 | |
| Other | 0.006566 | | 0.092666 | | 0.000227 | | 0.74573 | | 0.004502 | | 0.010244 | |

**S7. Two-tailed t-test analysis of changes in relative abundances of the dominant bacterial taxa in the respiratory microbiota between control and H9N2 AIV infected groups at different time points.** Two-tailed t-tests were used to assess how the relative abundances of bacterial taxa at phylum, class, order and family level change between control chickens and chickens challenged with H9N2 AIV, compared at the same time point tested. Samples of the respiratory microbiota were taken at days 2 (D2), 4 (D4) and 10 (D10) post-challenge in both groups. Only bacterial taxa that accounted for more than 1% of total sequences were included in the graphs. All other bacterial taxa were categorised as ‘Other’.

| **Bacterial Taxa** | **Days Post-Challenge** | | |
| --- | --- | --- | --- |
|  | **D2** | **D4** | **D10** |
| **Phylum Level** | | | |
| Actinobacteria | 0.101986 | 0.363521 | 0.02832 |
| Firmicutes | 0.031744 | 0.957259 | 0.461521 |
| Proteobacteria | 0.099331 | 0.584721 | 0.723955 |
| Other | 0.713574 | 0.121416 | 0.008364 |
| **Class Level** | | | |
| Actinobacteria | 0.102668 | 0.368062 | 0.023773 |
| Bacilli | 0.013956 | 0.96375 | 0.314503 |
| Clostridia | 0.1374 | 0.984636 | 0.459807 |
| Alphaproteobacteria | 0.085207 | 0.709262 | 0.255356 |
| Gammaproteobacteria | 0.147105 | 0.54864 | 0.505935 |
| Other | 0.980731 | 0.132289 | 0.079513 |
| **Order Level** | | | |
| Corynebacteriales | 0.016625 | 0.684078 | 0.004009 |
| Propionibacteriales | 0.036689 | 0.229746 | 0.01436 |
| Bacillales | 0.070597 | 0.449441 | 0.012673 |
| Lactobacillales | 0.00436 | 0.86831 | 0.059004 |
| Clostridiales | 0.1374 | 0.984636 | 0.459807 |
| Betaproteobacteriales | 0.082501 | 0.576468 | 0.353347 |
| Enterobacteriales | 0.233271 | 0.223211 | 0.050248 |
| Pseudomonadales | 0.951982 | 0.642295 | 0.045553 |
| Other | 0.195132 | 0.158532 | 0.083361 |
| **Family Level** | | | |
| Corynebacteriaceae | 0.020438 | 0.679493 | 0.004705 |
| Propionibacteriaceae | 0.046992 | 0.237754 | 0.016283 |
| Staphylococcaceae | 0.067708 | 0.43341 | 0.01497 |
| Lactobacillaceae | 0.004232 | 0.827726 | 0.042089 |
| Family XI | 0.336359 | 0.641122 | 0.103631 |
| Lachnospiraceae | 0.23259 | 0.358378 | 0.885705 |
| Ruminococcaceae | 0.168313 | 0.320158 | 0.841101 |
| Burkholderiaceae | 0.116916 | 0.56359 | 0.496488 |
| Enterobacteriaceae | 0.233271 | 0.223211 | 0.050248 |
| Moraxellaceae | 0.03933 | 0.92905 | 0.130859 |
| Pseudomonadaceae | 0.485736 | 0.097844 | 0.384672 |
| Other | 0.082999 | 0.169418 | 0.029697 |
