## Supplementary figures for "Alteration of the chicken upper respiratory microbiota, following H9N2 avian influenza virus infection": Supplementary_figures_Microbiota_ChrzastekK.docx

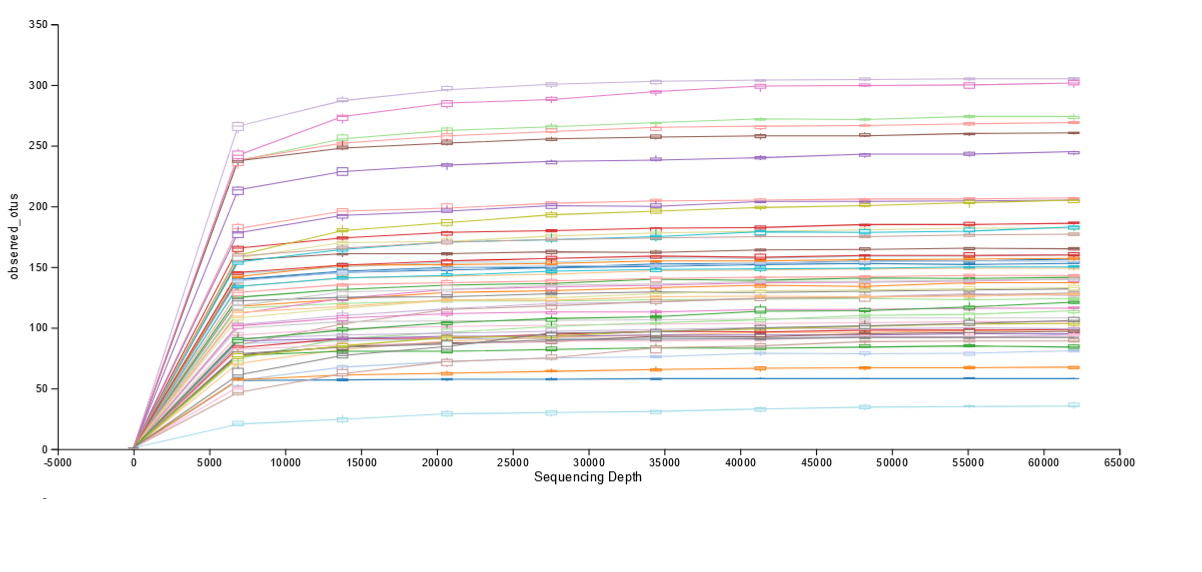

Suppl. Figure 1. Rarefaction curve for the observed number of OTUs. Rarefaction curve for the number of OTUs observed in each sample in both Experiment 1 and Experiment 2. Each colour represents individual sample. The OTUs were obtained from 16S rRNA gene sequencing of the V2-V3 region. All samples were normalised to 62,000 sequences per sample.

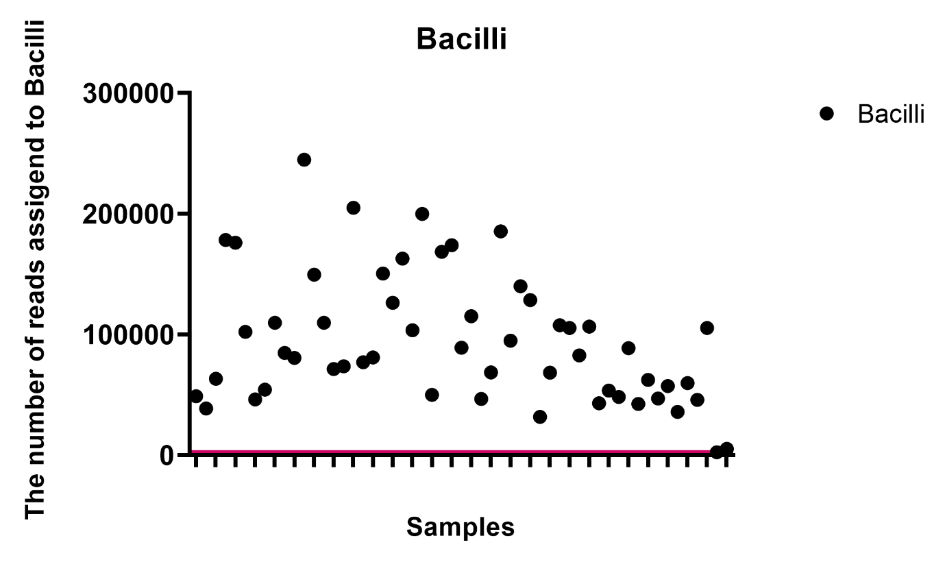

Suppl. Figure 2. The number of reads assigned to Bacilli per sample. The red line indicates the number of Bacilli reads found in the negative control that was used as a threshold.

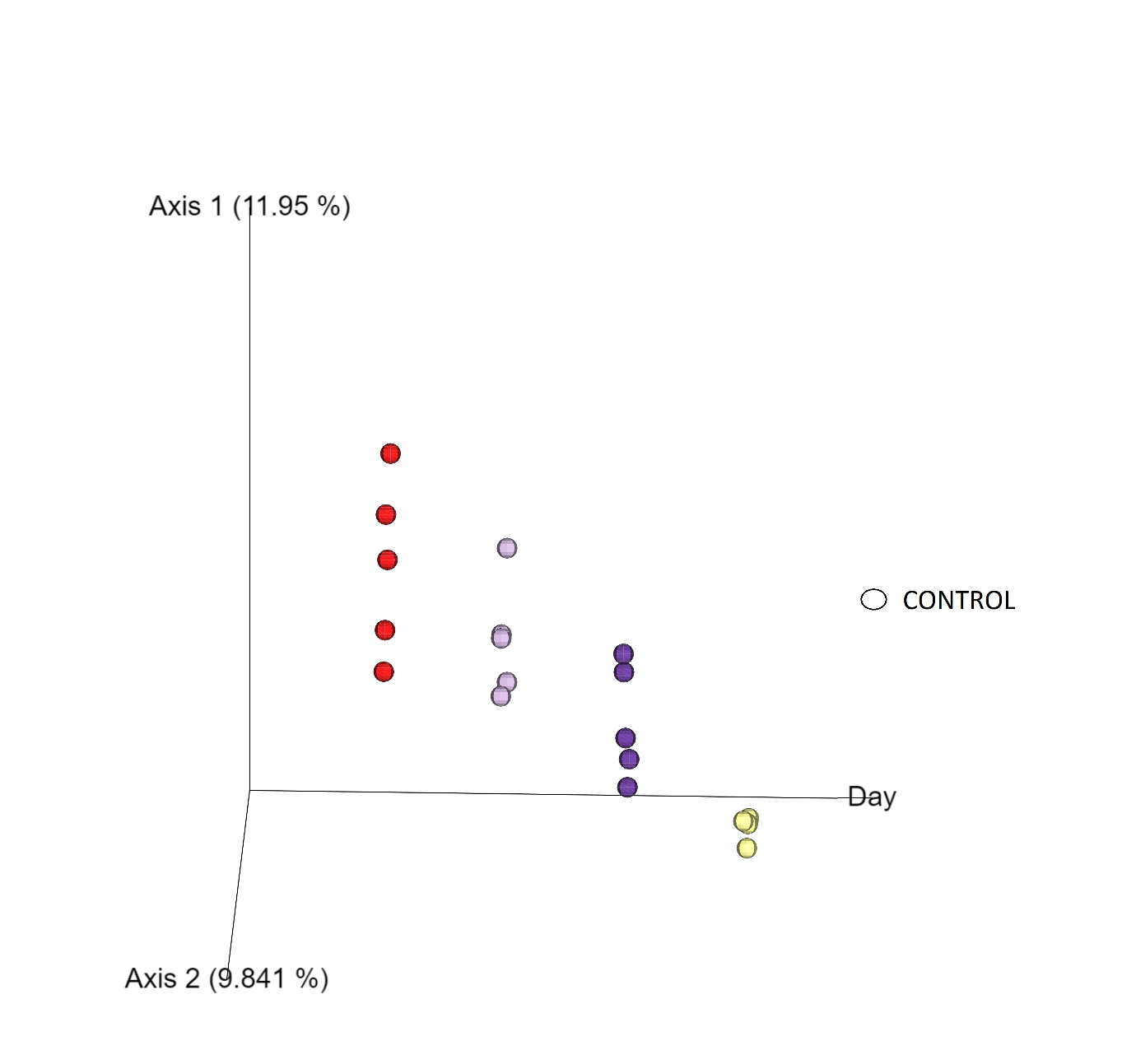

Suppl. Figure 3. A principal coordinate analysis (PCoA) plot of the healthy chicken upper respiratory track microbiome categorised by day using unweighted UniFrac distances. The samples were collected at days 7 (red), 14 (light violet), 21 (dark violet) and 28 (yellow) of age.

Suppl. Figure 4. Bacterial taxa that dominate the healthy URT microbiota during maturation. Mean relative abundances of bacterial (A) phylum (B) class (C) order (D) family are expressed as percentages. Samples of the respiratory microbiota were taken at days 7 (D7), 14 (D14), 21 (D21) and 28 (D28) of age. Only bacterial taxa that accounted for more than 1% of total sequences were included in the graphs. All other bacterial taxa were categorised as ‘Other’.

(C)

(D)

(A)

(B)

Relative Abundance (%)

|  | Actinobacteria_Actinobacteria_Corynebacteriales | Actinobacteria_Actinobacteria_Propionibacteriales | Firmicutes_Bacilli_Bacillales | Firmicutes_Bacilli_Lactobacillales | Firmicutes_Clostridia_Clostridiales | Proteobacteria_Gammaproteobacteria_Betaproteobacteriales | Proteobacteria_Gammaproteobacteria_Enterobacteriales | Proteobacteria_Gammaproteobacteria_Pseudomonadales | Other |
| --- | --- | --- | --- | --- | --- | --- | --- | --- | --- |
| D7 | 0.0919835 | 0.137153 | 1.07641 | 52.0164 | 29.3295 | 0.464388 | 14.6931 | 1.0887 | 1.10237 |
| D14 | 0.754679 | 0.592037 | 7.19923 | 32.297 | 21.5369 | 0.831767 | 29.1099 | 4.03522 | 3.64326 |
| D21 | 0.560118 | 0.710942 | 13.6934 | 16.4633 | 36.5136 | 1.52677 | 16.2445 | 9.96744 | 4.31986 |
| D28 | 0.19665 | 0.349458 | 4.87438 | 29.8788 | 48.7528 | 0.48166 | 11.8379 | 0.716424 | 2.91196 |

Relative Abundance (%)

|  | Actinobacteria_Actinobacteria_Corynebacteriales_Corynebacteriaceae | Actinobacteria_Actinobacteria_Propionibacteriales_Propionibacteriaceae | Firmicutes_Bacilli_Bacillales_Staphylococcaceae | Firmicutes_Bacilli_Lactobacillales_Lactobacillaceae | Firmicutes_Clostridia_Clostridiales_Family XI | Firmicutes_Clostridia_Clostridiales_Lachnospiraceae | Firmicutes_Clostridia_Clostridiales_Ruminococcaceae | Proteobacteria_Gammaproteobacteria_Betaproteobacteriales_Burkholderiaceae | Proteobacteria_Gammaproteobacteria_Enterobacteriales_Enterobacteriaceae | Proteobacteria_Gammaproteobacteria_Pseudomonadales_Moraxellaceae | Proteobacteria_Gammaproteobacteria_Pseudomonadales_Pseudomonadaceae | Other |
| --- | --- | --- | --- | --- | --- | --- | --- | --- | --- | --- | --- | --- |
| D7 | 0.0919835 | 0.120511 | 0.926796 | 50.7661 | 0.665511 | 22.329 | 5.98787 | 0.462102 | 14.6931 | 1.01973 | 0.0689682 | 2.86833 |
| D14 | 0.754679 | 0.557355 | 6.5537 | 30.2525 | 6.36131 | 12.9807 | 1.59021 | 0.831767 | 29.1099 | 3.30239 | 0.732829 | 6.9726 |
| D21 | 0.555945 | 0.610075 | 13.2269 | 15.561 | 0.681228 | 31.8234 | 3.49873 | 1.52185 | 16.2445 | 8.96999 | 0.997459 | 6.30875 |
| D28 | 0.19665 | 0.347308 | 4.09509 | 25.9305 | 0.165402 | 36.2244 | 4.84597 | 0.39223 | 11.8379 | 0.530389 | 0.186036 | 15.2482 |

Relative Abundance (%)

|  | Actinobacteria_Actinobacteria | Firmicutes_Bacilli | Firmicutes_Clostridia | Proteobacteria_Alphaproteobacteria | Proteobacteria_Gammaproteobacteria | Other |
| --- | --- | --- | --- | --- | --- | --- |
| D7 | 0.300882 | 53.0928 | 29.3295 | 0.0765524 | 16.3514 | 0.848893 |
| D14 | 1.94977 | 39.4962 | 21.5369 | 1.57868 | 34.4849 | 0.953541 |
| D21 | 2.33654 | 30.1567 | 36.5136 | 1.93722 | 28.1801 | 0.875823 |
| D28 | 1.06092 | 34.7532 | 48.7528 | 0.96503 | 13.3175 | 1.15053 |

Relative Abundance (%)

|  | Actinobacteria | Firmicutes | Proteobacteria | Other |
| --- | --- | --- | --- | --- |
| D7 | 0.336905 | 82.7829 | 16.4989 | 0.381264 |
| D14 | 1.95332 | 61.2329 | 36.0642 | 0.749518 |
| D21 | 2.39944 | 66.9591 | 30.2481 | 0.393344 |
| D28 | 1.14656 | 84.1694 | 14.3269 | 0.357089 |

Suppl. Figure 5. Bacterial taxa associated with H9N2 infection in the respiratory microbiota. Mean relative abundances of bacterial (A) phyla (B) class (C) order (D) family are expressed as percentages. Samples of the respiratory microbiota were taken at day 0 pre-challenge (D0) and at days 2 (D2), 4 (D4) and 10 (D10) post-challenge in both control chickens and chickens challenged with H9N2 AIV A/chicken/Pakistan/UDL01/08. Only bacterial taxa that accounted for more than 1% of total sequences were included in the graphs. All other bacterial taxa were categorised as ‘Other’.

(C)

(A)

(B)

(D)

Relative Abundance (%)

|  | Actinobacteria_Actinobacteria_Corynebacteriales | Actinobacteria_Actinobacteria_Propionibacteriales | Firmicutes_Bacilli_Bacillales | Firmicutes_Bacilli_Lactobacillales | Firmicutes_Clostridia_Clostridiales | Proteobacteria_Gammaproteobacteria_Betaproteobacteriales | Proteobacteria_Gammaproteobacteria_Enterobacteriales | Proteobacteria_Gammaproteobacteria_Pseudomonadales | Other |
| --- | --- | --- | --- | --- | --- | --- | --- | --- | --- |
| Control D0 | 1.71314 | 5.06717 | 12.6169 | 28.7028 | 17.0824 | 3.02744 | 25.2885 | 1.41233 | 5.08931 |
| Control D2 | 1.76927 | 2.00514 | 18.0267 | 27.5665 | 14.0726 | 4.31192 | 18.0909 | 4.57988 | 9.57704 |
| Infected D2 | 0.255602 | 0.451299 | 0.810413 | 81.9427 | 3.75287 | 0.97214 | 4.1035 | 4.88361 | 2.82784 |
| Control D4 | 1.81648 | 3.39315 | 11.4747 | 46.5841 | 8.59897 | 2.15621 | 10.7977 | 7.26623 | 7.91249 |
| Infected D4 | 1.30608 | 1.40858 | 7.08934 | 50.1823 | 8.52417 | 1.4769 | 22.1851 | 5.34823 | 2.4792 |
| Control D10 | 3.10298 | 3.59948 | 23.1747 | 17.0897 | 21.048 | 3.26022 | 7.36963 | 8.09838 | 13.257 |
| Infected D10 | 0.279429 | 0.984975 | 6.81316 | 47.4359 | 13.7975 | 2.0054 | 21.4234 | 2.68722 | 4.57303 |

Relative Abundance (%)

|  | Actinobacteria_Actinobacteria_Corynebacteriales_Corynebacteriaceae | Actinobacteria_Actinobacteria_Propionibacteriales_Propionibacteriaceae | Firmicutes_Bacilli_Bacillales_Staphylococcaceae | Firmicutes_Bacilli_Lactobacillales_Lactobacillaceae | Firmicutes_Clostridia_Clostridiales_Family XI | Firmicutes_Clostridia_Clostridiales_Lachnospiraceae | Firmicutes_Clostridia_Clostridiales_Ruminococcaceae | Proteobacteria_Gammaproteobacteria_Betaproteobacteriales_Burkholderiaceae | Proteobacteria_Gammaproteobacteria_Enterobacteriales_Enterobacteriaceae | Proteobacteria_Gammaproteobacteria_Pseudomonadales_Moraxellaceae | Proteobacteria_Gammaproteobacteria_Pseudomonadales_Pseudomonadaceae | Other |
| --- | --- | --- | --- | --- | --- | --- | --- | --- | --- | --- | --- | --- |
| Control D0 | 1.71314 | 4.93185 | 12.4846 | 26.8592 | 0.92125 | 12.6225 | 2.70681 | 2.11437 | 25.2885 | 0.816868 | 0.595466 | 8.94544 |
| Control D2 | 1.67098 | 1.8207 | 17.7694 | 25.977 | 1.80747 | 9.37082 | 2.2192 | 3.88971 | 18.0909 | 3.40936 | 1.17052 | 12.8039 |
| Infected D2 | 0.229817 | 0.451299 | 0.728522 | 81.7854 | 0.772648 | 2.72789 | 0.247102 | 0.969173 | 4.1035 | 0.423155 | 4.46046 | 3.10101 |
| Control D4 | 1.81648 | 3.28102 | 11.3042 | 44.4439 | 4.0281 | 3.17278 | 1.17675 | 2.09316 | 10.7977 | 5.2856 | 1.98063 | 10.6197 |
| Infected D4 | 1.30008 | 1.37989 | 6.85823 | 49.3697 | 2.72045 | 4.83275 | 0.732185 | 1.38833 | 22.1851 | 4.97528 | 0.372952 | 3.885 |
| Control D10 | 2.6515 | 3.38523 | 22.4753 | 12.6109 | 5.76217 | 10.7944 | 3.57234 | 2.88077 | 7.36963 | 6.05486 | 2.04352 | 20.3994 |
| Infected D10 | 0.279429 | 0.961538 | 6.63145 | 47.0035 | 0.838783 | 9.74206 | 3.08544 | 2.00097 | 21.4234 | 1.90621 | 0.781006 | 5.3463 |

Relative Abundance (%)

|  | Actinobacteria | Firmicutes | Proteobacteria | Other |
| --- | --- | --- | --- | --- |
| Control D0 | 7.8459 | 58.606 | 32.5499 | 0.998197 |
| Control D2 | 7.97205 | 59.8173 | 31.1418 | 1.06884 |
| Infected D2 | 1.26666 | 86.605 | 11.3763 | 0.752009 |
| Control D4 | 6.3009 | 66.7978 | 23.9749 | 2.9264 |
| Infected D4 | 3.33841 | 66.0181 | 30.3466 | 0.296854 |
| Control D10 | 9.10884 | 61.7114 | 25.3808 | 3.79896 |
| Infected D10 | 2.55655 | 69.0855 | 28.1355 | 0.222433 |

Relative Abundance (%)

|  | Actinobacteria_Actinobacteria | Firmicutes_Bacilli | Firmicutes_Clostridia | Proteobacteria_Alphaproteobacteria | Proteobacteria_Gammaproteobacteria | Other |
| --- | --- | --- | --- | --- | --- | --- |
| Control D0 | 7.75959 | 41.3197 | 17.0824 | 1.50605 | 31.0349 | 1.2974 |
| Control D2 | 7.89198 | 45.5932 | 14.0726 | 3.32988 | 27.7908 | 1.32152 |
| Infected D2 | 1.26064 | 82.7531 | 3.75287 | 0.972344 | 10.404 | 0.857064 |
| Control D4 | 6.265 | 58.0588 | 8.59897 | 1.31945 | 22.6393 | 3.11846 |
| Infected D4 | 3.33589 | 57.2717 | 8.52417 | 1.04321 | 29.3034 | 0.521632 |
| Control D10 | 8.26924 | 40.2644 | 21.048 | 3.22933 | 21.5689 | 5.62023 |
| Infected D10 | 2.5508 | 54.2491 | 13.7975 | 1.42237 | 26.4759 | 1.50435 |

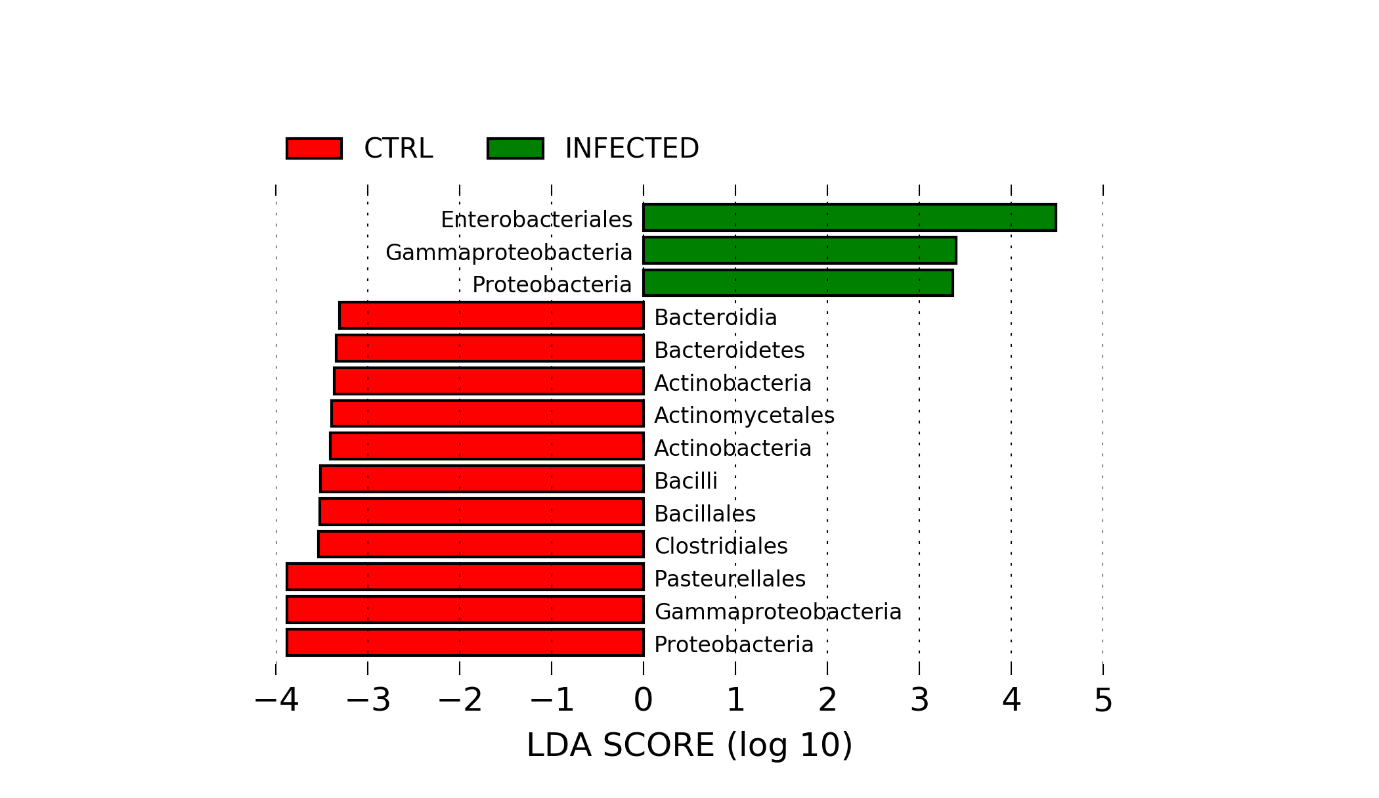

Suppl. Figure 6. Linear discriminant analysis Efect Size (LEfSe) identifying differences between H9N2 AIV infected chickens and controls at day 4 post-challenge. Control birds are represented by red whereas infected birds are represented by green colour. The birds were challenged with 10^5^ PFU/ml recombinant H9N2 A/chicken/Pakistan/UDL01/01. Kruskal-Wallis and Wilcoxon tests were applied at alpha value 0.05. LDA scores (log10) above 2.0 and p < 0.05 are shown.

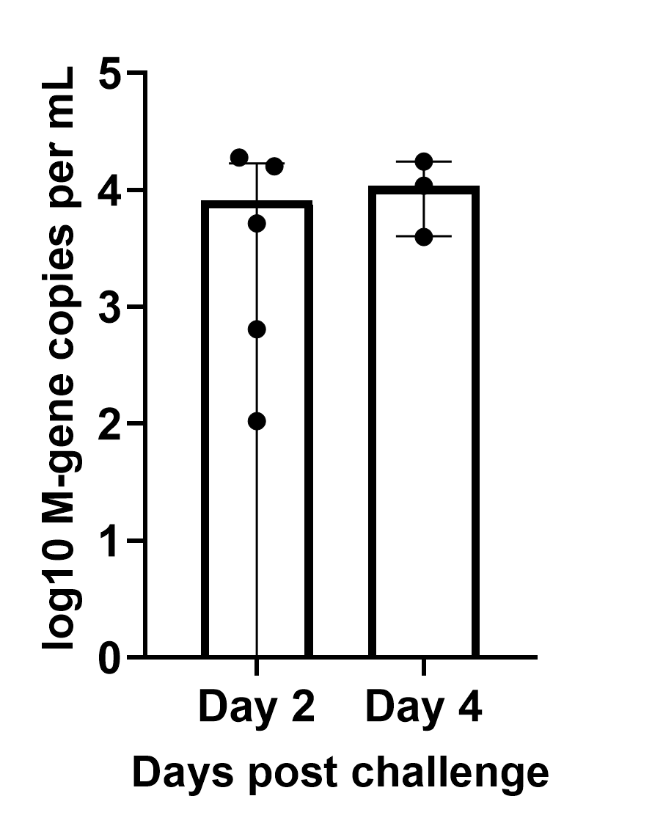

Suppl. Figure 7. The number of copies per ml of the influenza A virus matrix gene (M-gene) detected in trachea samples taken from chickens at days 2 and 4 post-challenge with H9N2 AIV A/chicken/Pakistan/UDL01/08 by qRT-PCR. The number of M-gene copies per ml is expressed as Log10.
